# Immune evasion impacts the selective landscape of driver genes during tumorigenesis

**DOI:** 10.1101/2022.06.20.496910

**Authors:** Lucie Gourmet, Andrea Sottoriva, Maria Secrier, Luis Zapata

## Abstract

Carcinogenesis is an evolutionary process fueled by the interplay of somatic mutations and the local microenvironment. In recent years, hundreds of cancer related genes have been discovered using cancer cohorts. However, these cohorts are heterogenous mixtures of different molecular phenotypes, which hampers the identification of driver genes associated to a specific cancer hallmark or microenvironment. Here, we compared the landscape of positively selected somatic mutations in immune-escaped (escape+) versus non-escaped (escape-) tumors. We applied the ratio of non-synonymous to synonymous mutations (dN/dS) to 9896 individuals from 31 primary tumor tissues from the Cancer Genome Atlas (TCGA) separated by escape status. Altogether, we found 85 driver genes, including 27 and 16 novel driver genes in escape- and escape+ tumors, respectively. Overall, driver dN/dS of escape+ tumors (dN/dS=1.23) was significantly lower and closer to neutrality than driver dN/dS of escape-tumors (dN/dS=1.62), suggesting a relaxation of positive selection in driver genes, a relaxation of negative selection on immunogenic driver sites, or a combination of both fueled by immune escape. We also found that the proportion of unique sites mutated in escape+ tumors is almost double than in escape-tumors, and that immune evasion allows for a more diverse repertoire of mutational signatures. We also identified that strong immunoediting in the absence of escape leads to a better overall survival in tumors enriched by an inflamed phenotype. Ultimately, our findings reveal differences in the evolutionary strategies used by cancer cells to establish tumorigenesis and highlight the need for better patient stratification to develop tailored treatments based on molecular targets.

## Introduction

Cancer is a highly prevalent disease defined by genetic instability and the accumulation of mutations. In the 1970s, Peter Nowell described cancer as a multistage process subject to different selective pressures [1]. Somatic mutations in driver genes promote cancer by providing cells with a proliferative advantage. Over the last decade, several efforts have focused on the discovery of novel cancer drivers [2]. However, the landscape of driver genes remains incomplete as an increasing number of driver genes are identified as more tumors are sequenced. In addition to driver mutations, passenger mutations occur randomly and have a neutral role regarding tumourigenesis. Not only are passenger mutations more frequently observed than drivers, but they also provide a record of traces left by mutagenic exposure [3]. Meanwhile, negative selection eliminates cells carrying deleterious or antigenic mutations [4]. However, unlike positive selection, the role and extent of negative selection during tumor evolution is controversial.

To detect genes under selection, the ratio of nonsynonymous to synonymous mutations (dN/dS) has been a widely used metric in comparative genomics. Genes under positive selection harbor more protein-altering mutations than expected by chance (dN/dS > 1), unlike genes under negative selection that are depleted of such mutations (dN/dS < 1). The mutational background is important as the combination of some genetic variations, or mutually exclusive mutations, can lead to cell death. This also implies that the perturbation of pairs of genes can result in synthetic lethality, which can be exploited to develop new treatments. To identify the full landscape of driver genes, we need to consider mutational processes [5] and molecular phenotypes associated to cancer progression.

Cancer immunoediting is an evolutionary process that selects for clones with low immunogenicity or the ability to escape immune recognition. Cancer camouflage can occur through the depletion of immunogenic variants, downregulation of tumor antigen expression, or an overexpression of checkpoint proteins, such as PD1 and CTLA4 [7]. Somatic mutations can lead to new peptides, known as neoantigens, which are processed by the Major Histocompatibility Complex (MHC) and displayed on the surface of the tumor for the immune system to recognize. Upon recognition, these cancer cells can be eliminated by various immune cells, including cytotoxic T cells or natural killer cells. The mutational landscape can be linked to survival as clonal neoantigens were found to affect the sensitivity to immune checkpoint blockade [6].

However, the tumor microenvironment can also promote cancer progression (e.g. cancer associated fibroblasts are known to secrete growth factors, and macrophages can support angiogenesis) [7,8]. This demonstrates that the tumor microenvironment can also be immunosuppressive and favorable to carcinogenesis. In fact, the microenvironment has also been linked to metastasis since it can serve as a preme-tastatic niche [7]. A study comparing metastatic and primary tumors demonstrated that tumors which had metastasized contained fewer cytotoxic T cells [8]. The microenvironment differs between the primary and different metastatic sites, as it has been demonstrated in ovarian and breast cancer [9, 10]. This is likely due to immunoediting, as unedited or non-immunogenic clones were the only ones to survive in a study of 31 colorectal metastases with exceptionally long survival [11]. However, the differential selective landscape between immune escape+ and escape-tumors has not yet been explored.

Here, we stratified patients into two cohorts based on somatic mutations in immune-related genes and compared their selective landscape. We applied dN/dS in these two cohorts and found 85 driver genes that could be linked to the presence or absence of an active immune selective process. We demonstrated that several known cancer genes accumulate mutations in specific mutational hotspots only in escape-groups, suggesting the active role of immunoediting on shaping the distribution and frequency of driver events. Finally, immune escaped individuals have worse overall prognosis in immune inflamed tumors, possibly linked to the plasticity required to colonize new niches.

## Results

### Immune escape leads to neutral-like evolutionary dynamics measured by dN/dS

To determine the impact of immune evasion in the selective landscape of tumorigenesis, we obtained a catalog of 88 genes involved in the antigen presenting machinery or previously associated to immune evasion [12]. We classified 9896 TCGA patients from 31 different cancer subtypes into escaped (escape+) and non-escaped (escape-) cohorts based on the presence of a mutation in one of these genes (Fig 1). These resulted into 2089 escape+ individuals with an average tumor mutation burden per patient (TMB) of 426 - over 4 times higher than the average TMB for the 7087 escape-patients (95 mutations per individual, Supp Table 1). Specifically, we observed that the average number of mutations per individual in escape+ was 4.51 times higher for missense, and 3.95 times higher for truncating mutations compared to escape-tumors (Supp Table 1). Other mutation types, such as splicing and synonymous events, were also higher in escape+ (Supp Fig. 1).

**Figure 1.**
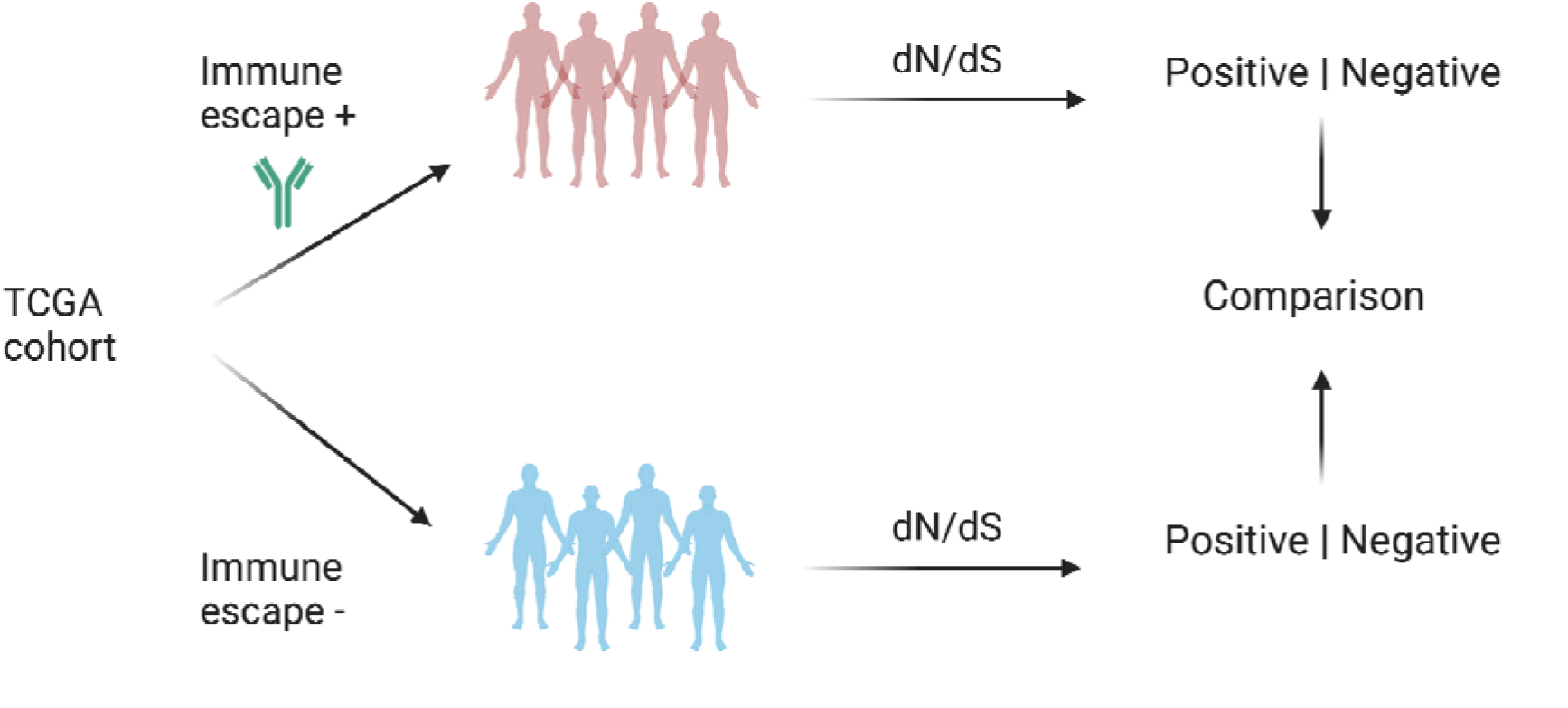
9896 tumors across 31 cancer subtypes from TCGA were classified into escape+ and escape-based on mutations presented in the antigen presenting machinery. These two cohorts were then analysed to detect genes under significant selection using dN/dS. Genes under significant selection can be used as molecular targets to improve cancer treatment strategies.

We then calculated dN/dS on missense and truncating mutations for the escape+ and escape-cohorts. We calculated global dN/dS using all genes and driver dN/dS using a set of 366 known driver genes for each tumor type and the whole cohort [2]. At the pancancer level, we observed that global dN/dS (Escape+: 1.05, CI=[1.041:1.051], Escape-:1.07, CI=[1.060:1.072], Supplementary Fig 2) and driver dN/dS (Fig 2A, Escape+:1.223, CI=[1.189:1.257], Escape-:1.619, CI=[1.57:1.669]) in escape+ were lower and closer to neutrality compared to escape-, suggesting that each group had evolved under different selective pressures.

**Figure 2.**
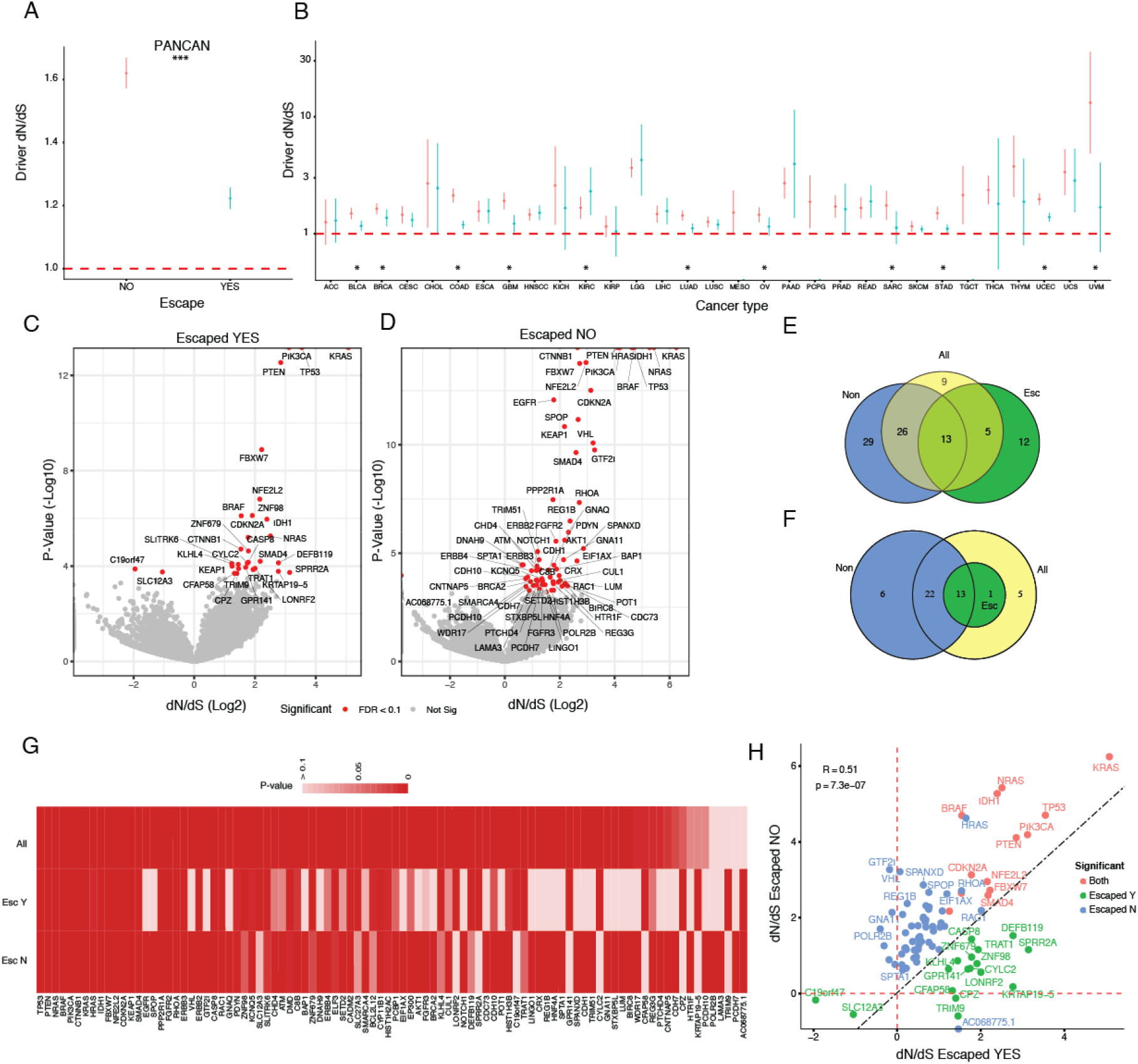
Landscape of selection in escape+ and escape-tumors. A) Overall driver dN/dS for escape+ and escape-tumors across all TCGA tumor types. B) Overall driver dN/dS for 31 cancer subtypes separated by escape status (red – escaped WT, blue – escaped Mut). Volcano plot for gene-level dN/dS using missense mutations versus P-value. Venn diagram showing the number of significant driver genes (Q-Value < 0.1) using missense mutations considering E) all genes or F) restricted to known driver genes when separating by escape group or with all samples together. G) List of significant driver genes using missense mutations in at least one group. H) dN/dS value for significant genes in escaped WT versus escaped Mut groups.

When looking at each tumor subtype, global dN/dS was significantly higher than one in 25 out of 31 escape- and in 19 out of 31 escape+ tumors (Supplementary Fig 3). Driver dN/dS was significantly higher than one in 29/31 escape-tumors compared to 20/28 escape+ tumors. Overall, global dN/dS was similar between escape- and escape+, with a majority (21 out of 31) of escape-having higher global dN/dS compared to escape+. In ACC, GBM and UVM, dN/dS was significantly higher in the escape-group, whereas in KIRP, dN/dS was significantly higher in the escape+ group. Driver dN/dS was higher in all, and significantly higher in 11, escape-tumors compared to escape+ (Fig 2B). Overall, immune escaped tumors have close to neutral dN/dS suggesting that immune evasion relaxes positive selection of driver events, which is possibly linked to a lower selective advantage needed to initiate tumorigenesis in the absence of functional immune surveillance.

Next, we hypothesized that if global and driver dN/dS are different between escaped and non-escaped groups, the landscape of driver genes would also be different. Thus, to uncover novel driver genes in escape+ and escape-patients, we calculated dN/dS for missense and truncating mutations in all genes. For missense mutations, we found 30 and 68 significant genes in escape+ (Fig 2C) and escape- (Fig 2D) tumors, respectively. 17 out of 30 were specific to escape+ and 55 out of 68 were specific to escape-, with 13 driver genes common to both groups. For truncating events, we found 64 and 41 driver genes for escape- and escape+ tumors, respectively, with 33/64 specific to escape-, 10/41 specific to escape+ and 31 common to both (Fig S4). To determine whether stratifying patients into molecular subgroups revealed novel driver genes, we calculated dN/dS using all patients together. From the set of 68 significant genes in escape-, 29 of them were missed when calculating dN/dS using all patients (Fig 2E). Similarly, 12 driver genes were only found in the escape+ group, highlighting the impact of mixing patients with different evolutionary paths into cohort analysis for the discovery of cancer driver genes. Moreover, if we restrict this analysis to only known driver genes, we still observed 6 genes under significant selection only in the non-escaped group, which were missed when combined with escape+ patients (Fig 2F). Importantly, when combining all patients to predict driver genes, the majority still has a significant P-value (89 out of 94 genes), but lost significance after multiple testing correction (Fig 2G), hence the importance of properly stratifying groups.

Among 55 genes under significant positive selection in the escape-group, the majority (41/55) was evolving neutrally in the escape+ group despite having a similar number of mutations (Fig S5). We then compared dN/dS in significant genes in escape+, escape– or common. We found that in escape+ significant genes, most of the genes have also signals of positive selection in the escape-group but with values closer to neutrality (Fig S6). Furthermore, we also found two genes under significant negative selection specifically in the escape+ group: SLC12A3 and C19orf47. In the common genes, there was an overall higher dN/dS in the non-escaped group (9 out of 13) and all were previously known drivers (Fig S7). The gene-level driver dN/dS value of escape-versus escape+ was significantly correlated in the pancancer analysis. However, in every case, there were genes under strong positive selection in one group and not in the other, thereby validating the fact that opposing evolutionary pressures shape differently the landscape of driver genes. Among genes under strong positive selection in escape-but not in escape+ genes, we found GTF2I, VHL, REG1B, SPANXD, in contrast to other genes such as CPZ, CRTAP19-5, CFAP58, that are under strong positive selection in the escape+ but neutrally evolving in the escape-.

These results confirm our previous analysis of higher driver dN/dS in escape-pancancer and across cancer types, suggesting that immune evasion relaxes the selective advantage needed for tumorigenesis. The fact that we did not find a higher number of driver genes in escape+ suggests that the relaxation of selection comes as a fitness trade-off of common driver genes in selected spots which can be favored when the immune system is absent as they are less immunogenic.

### Mutations are evenly distributed across driver genes in immune-escaped patients

To determine the influence of the immune system on the landscape of driver mutations, we explored whether mutations occurred more often at specific sites in escape+ compared with escape-patients. We first compared two genes that were significant drivers in both scenarios, IDH1 and KRAS. We observed that IDH1 is under positive selection in both groups but with a significant higher dN/dS in the escape-group (dNdS of 31 versus dNdS of 3, Fig 3A). In contrast, KRAS also has a higher dN/dS in the escape-compared to the escape+ group, but ultimately, it is not significantly different (dNdS of 75 versus dNdS of 35, Fig 3B). We found the common hotspots for IDH1 (Fig 3C, R132) and KRAS (Fig 3D, G12) as the most frequently mutated position in both groups. However, in escape-, a larger burden of mutations was concentrated in the hotspot while the number of unique sites mutated remained similar (Fig 3E, chi-2 p-val = 3.6e-12, Fig 3F chi-2 p-val 0.01). We then performed the same test on all significant common driver genes together (Pandriver) and found that the majority of cancer drivers harbor mutations at specific sites in escape-patients (hotspots), while mutations where evenly distributed across the gene in escape+ patients (Fig 3G, Pandriver p-value= 2.49e−53). This was also the case for other known driver genes such as BRAF (p-value =3.01E-08), TP53 (p-value =3.41E-11), EGFR (p-value = 0.00877) and GNA11 (p-value =0.00689), among others (Fig 3G). Interestingly, recent work by Hoyos et al demonstrated that TP53 hotspots are bound to fitness trade-offs between oncogenicity and immunogenicity, thereby suggesting that these hotspot mutations have reduced antigen presentation and therefore should not be present in escaped+ tumors. This ultimately validates our finding that escaped-tumors require an increase in intrinsic fitness to overcome the immune system.

**Figure 3.**
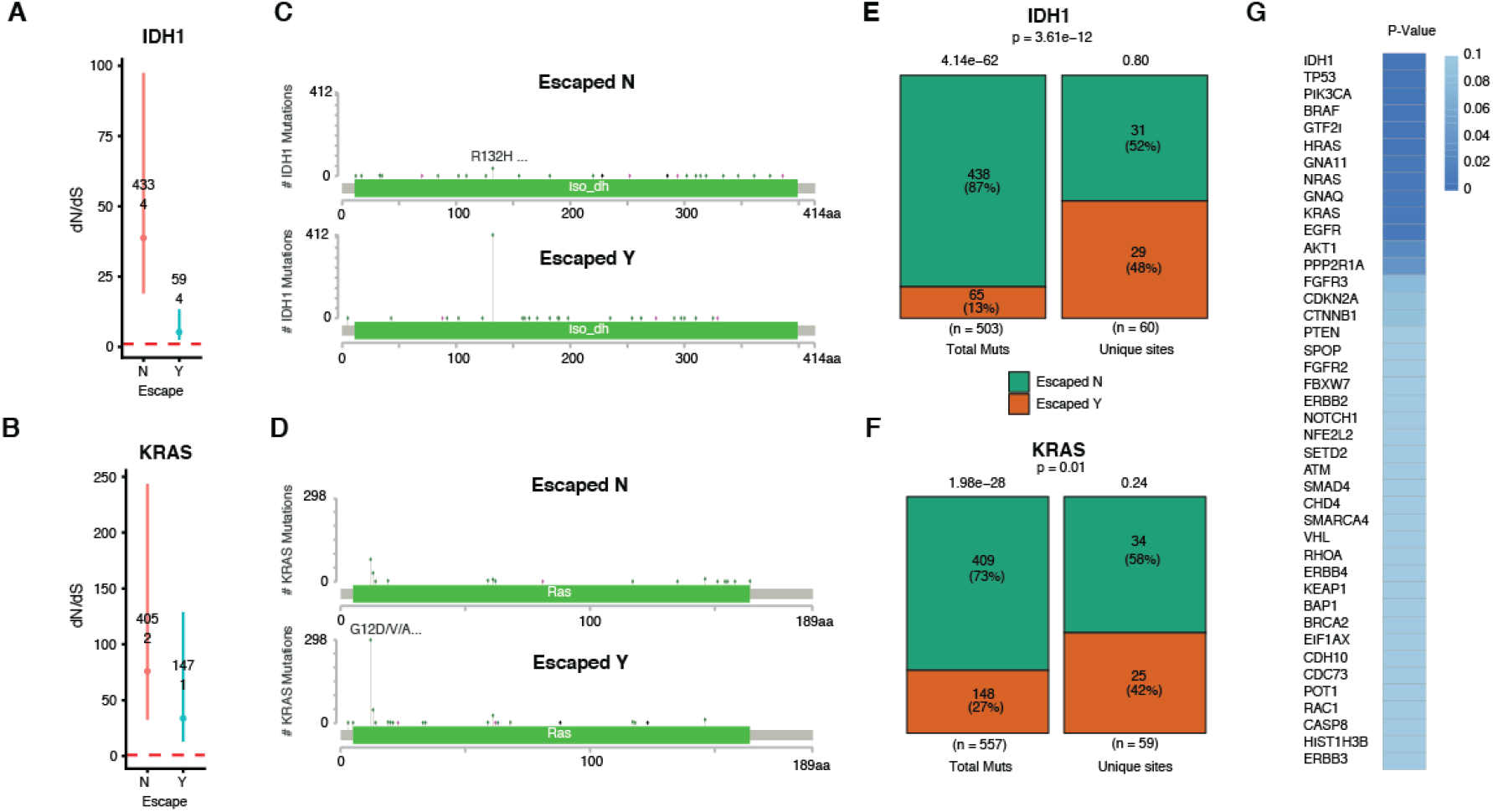
Non-random mutational distribution in non-escaped tumors. dN/dS for A) IDH1 and B) KRAS in escape-and escape+ groups including number of nonsynonymous (upper number) and synonymous (lower number) mutations. Lolliplots for mutations in escape- and escape+ tumors for C) IDH1 and D) KRAS. Chi-square test comparing the number of mutations and the number of unique mutated sites for E) IDH1 and F) KRAS. G) Chi-2 P-value for significant known driver genes comparing the number of mutations versus the number of sites in both groups.

To test whether the larger number of patients in the escape-group bias the results, we repeated our analysis by downsampling escape-patients to match escape+ patients. We found that escape+ patients had significantly more unique sites mutated in IDH1 and BRAF genes, supporting the hypothesis that, for these genes, immune selection restricts the availability of driver sites, thereby forcing the accumulation of mutations in hotspots with less immunogenic capacity.

### Mutational signatures associated to immune evasion in cancer

To determine the mutational signatures associated to immune evasion we run deconstructSigs on each cohort separated. We found a different profile in the trinucleotide substitutions between escaped+ (Fig 4A) and escaped-(Fig 4B) tumors, especially in sites associated to C>A substitutions in a TCT context, and C>T substitutions in a TCA or TCC context.

**Figure 4.**
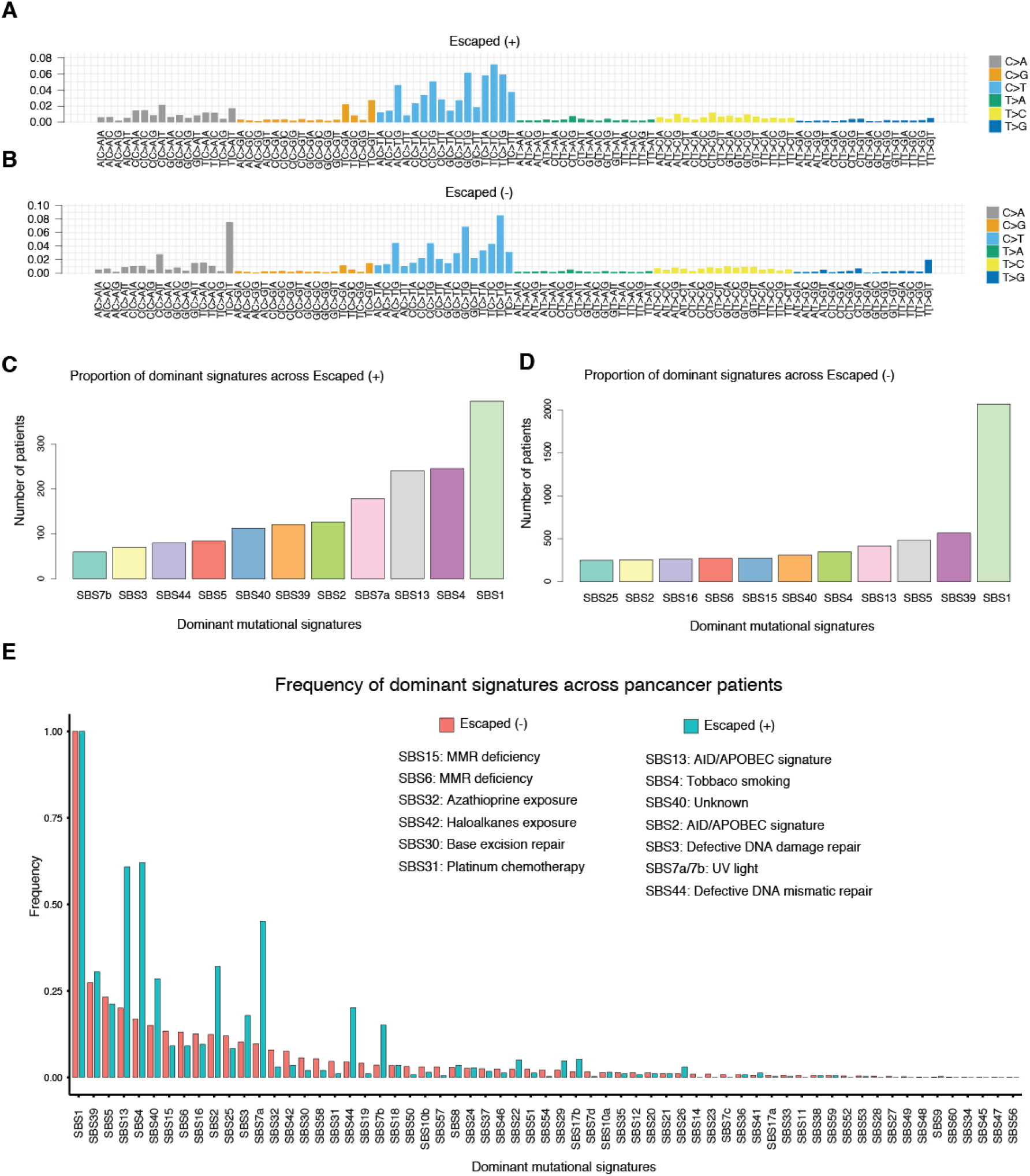
Mutational signatures associated to immune evasion. Proportion of mutation substitution in 96 trinucleotide contexts for A) escaped+ and B) escaped-tumor cohorts. Dominant signatures per patient in C) escaped+ and D) escaped-tumor cohorts. E) Distribution of frequency of the top 60 signatures in escaped-tumors and their distribution in escaped+ tumors.

We then tested whether immune escape allows a broader repertoire of mutational signatures to occur by comparing the most dominant signatures in escaped and non-escaped individuals. Despite in both groups the most dominant signature was SBS1, we observed a larger fraction of individuals harboring a different than SBS1 signature in the escape+ (Fig 4C) compared to the escape-group (Fig 4D). The top signatures for the escape+ group were SBS4 (smoking), SBS13 (APOBEC), SBS7a (UV exposure) and SBS2 (APOBEC). In contrast, the signatures in the pancancer escape-followed a flatter distribution with SBS39 and SBS5 being the second and third more frequent signatures. We next compared the top differential signatures between escaped+ and escaped-. Interestingly, we found that immune evasion was associated to APOBEC, tobacco and UV light signatures, while non-escaped tumors harbors signatures associated to mismatch repair deficiency and to various chemical exposures (Fig 4E).

### Immune inflammation leads to better prognosis in the absence of immune escape

To determine whether classifying tumors into immune categories can reveal different prognostic value between immune escaped and non-escaped tumors, we classified patients into 6 categories previously defined by Thorsson et al[17]. We found that the only category where there was a significant difference between groups was C3, which was characterized by an inflammatory signature. Immune-escaped tumors had a worst prognosis compared to non-escaped tumors (Fig 5), indicating that strong immunoediting in the absence of escape events leads to better survival. An observation which was recently demonstrated in long-term survivors of pancreatic cancer (Luksza et al 2022), which have stronger immunoediting signals compared to short-term survivors, characterized by weak immunoediting. Interestingly, one of the tumor types enriched in the C3 group is pancreatic adenocarcinoma.

**Figure 5.**
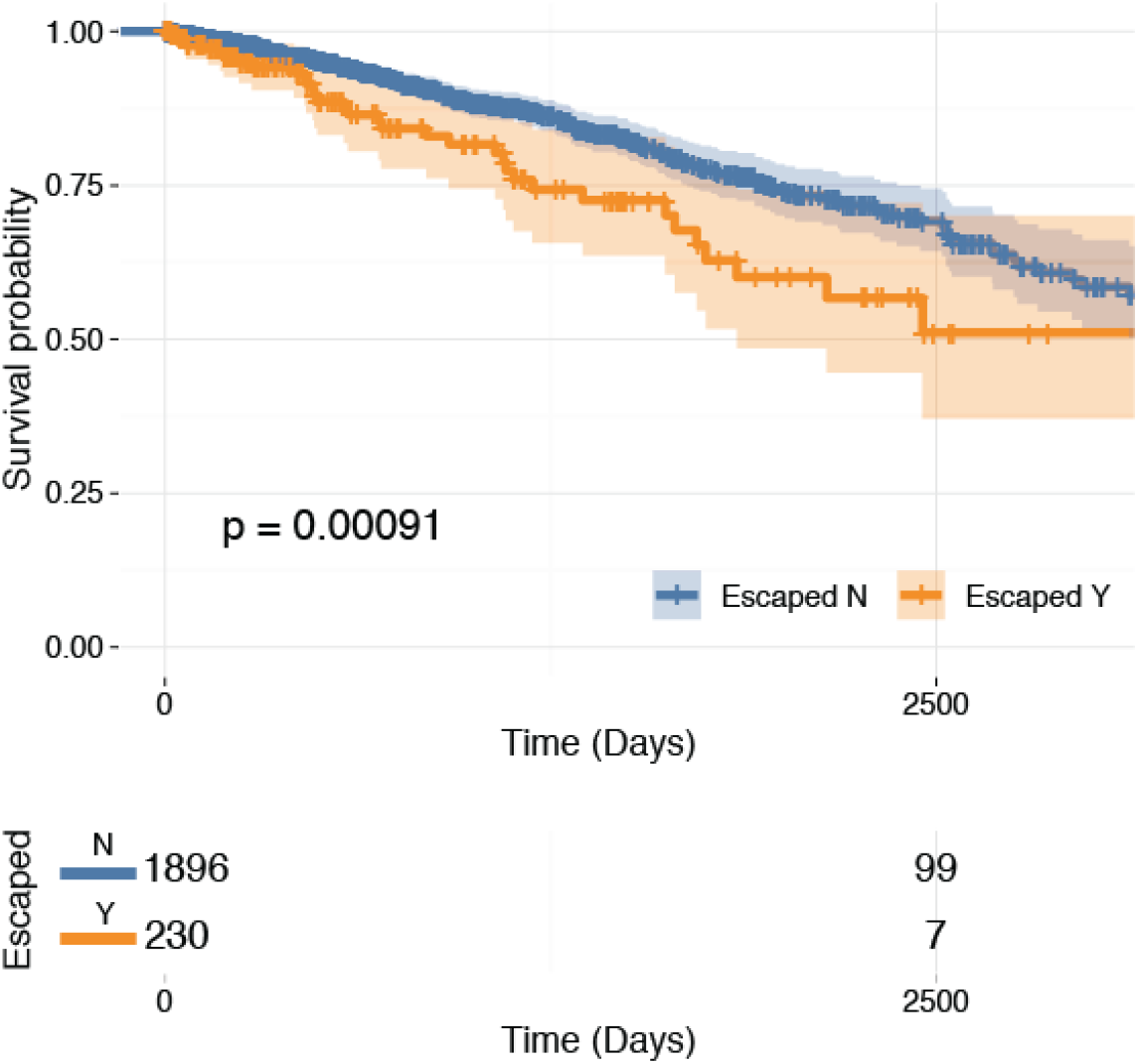
Overall survival for TCGA patients in immune category C3, characterized by an inflammatory signal.

Overall, we demonstrated key differences in the driver gene landscapes of escaped versus non-escaped tumors in primary tumor data by using dN/dS analysis. We observed that immune escape impacts the distribution of mutations within sites in known driver genes, especially in hotspots associated to lower neoantigen presentation. We found that mutational signatures are also associated to immune evasion mechanisms and that, by considering these two outcomes, we could significantly separate prognosis of primary tumors undergoing immune evasion in immune-inflamed tumors. Ultimately, our work highlights the importance of immune evasion as a hallmark of cancer and its effect as a compensatory mechanism when negative selection forces, such as the one exerted by the immune system, are in action.

## Discussion

In this study, we compared the selective landscape of immune escaped versus non-escaped tumors using the ratio of nonsynonymous to synonymous mutations, dN/dS. Using a cohort stratification based on the mutational status of immune-related genes, we discovered specific cancer drivers for each category and revealed new driver genes across tumor types in TCGA. We found that escaped tumors evolve more neutrally than non-escaped tumors when using dN/dS of all genes and of known driver genes as a metric of selection. By analyzing the distribution of mutations at unique sites, we identified that mutations are more evenly distributed across driver genes in escape+ patients. This result suggests that the relaxation of immune selection on driver sites leads to a dilution effect on dN/dS in common driver genes. Interestingly, a recent model explains the presence of hotspots events in driver genes in terms of a trade-off fitness advantage of oncogenicity versus immunogenicity, possibly confirming the differences we observe in the distribution of mutations between the escaped and non-escaped cohorts [15]. As we could expect from our findings with the distribution of mutations, the proportion of mutational signatures also varies between escape+ and escape-. Mutational signatures associated to immune evasion were associated to APOBEC and tobacco smoke while the signatures associated to non-escaped were related to mismatch repair and chemical exposure. Overall, we see that our stratification strategy reveals new potential target genes and a difference in selection. Our study sheds light on the evolution of the mutational landscape regarding the occurrence of mutations and mutational signatures. It also informs us on the prognosis of patients depending on their mutational background.

The importance of the interplay between the immune system and cancer cells has been widely studied from a neoantigen and MHC-class I perspective [12, 13]. A study by Grasso et al observed the role of genetic mechanisms of immune evasion in colorectal cancer [14]. They focused on the type of genetic instability (e.g. microsatellite) rather than on mutations linked to an escape status. In our work, we could identify a prognostic value of the escape status specifically in immune inflamed patients. A result in line with a recent analysis of pancreatic cancer patients, where Łuksza et al demonstrated that long-term survivors were subjected to strong immune editing, developed genetically less heterogeneous recurrent tumors with fewer neoantigens [16]. These observations highlight the need for a better understanding of the interaction between immune evasion, selection and clinical outcomes. Our study is the first to propose the stratification of immune evasion, or a surrogate of absence of immune-editing, as a prognostic biomarker during cancer evolution.

## Acknowledgements

LZ is supported by the ICR as a research fellow. A.S. is supported by the Wellcome Trust (202778/B/16/Z) and Cancer Research UK (A22909). We acknowledge funding from the National Institute of Health (NCI U54 CA217376) to A.S. This work was also supported by a Wellcome Trust award to the Centre for Evolution and Cancer (105104/Z/14/Z).

## Author contributions

LG performed all analysis and wrote the first draft of the paper. LZ conceived, designed, and supervised all analysis. MS and AS supervised the project. LZ and LG wrote the final version of the manuscript with the help of all the authors.

## Competing interests

All authors declare no financial/non-financial competing interest.

## Online Methods

### Dataset

We used The Cancer Genome Atlas (TCGA) project through the Genomics Data Commons (GDC) Portal (https://portal.gdc.cancer.gov/) to study 31 cancer types (downloaded in 13/10/2021). We included in the analysis: adrenocortical carcinoma (ACC, n=92 patients), bladder urothelial carcinoma (BLCA, n=411), breast cancer (BRCA, n=983), cervical cancer (CESC, n=288), Cholangiocarcinoma (CHOL, n=50), colorectal cancer (COAD n=390 and READ n=134, merged as CRC, n=524), esophageal cancer (ESCA, n=184), glioblastoma (GBM, n=390), head and neck squamous cell carcinoma (HNSC, n=506), Kidney Chromophobe (KICH, n=66), clear cell renal cell carcinoma (KIRC, n=336), papillary renal cell carcinoma (KIRP, n=281), low grade glioma (LGG, n=506), hepatocellular carcinoma (LIHC, n=364), lung adenocarcinoma (LUAD, n=567), lung squamous cell carcinoma (LUSC, n=490), Mesothelioma (MESO, n=81), ovarian cancer (OV, n=436), pancreatic cancer (PAAD, n=175), pheochromocytoma and paraganglioma (PCPG, n=179), prostate adenocarcinoma (PRAD, n=493), sarcoma (SARC, n=237), Skin Cutaneous Melanoma (SKCM, n=452), Stomach Adenocarcinoma (STAD, n=431, Testicular Germ Cell Tumors (TGCT, n=144), Thymoma (THYM, n=123), Thyroid carcinoma (THCA, n=492), Uterine Carcinosarcoma (UCS, n=56), Uterine Corpus Endometrial Carcinoma (UCEC, n=479) and Uveal Melanoma (UVM, n=80). This sums up to a pancancer cohort of 9896 patients.

For the immune category stratification, we used the categories C1 to C6 from Thorsson et al 2018 [18]. These are Wound Healing (C1), IFN-γ Dominant (C2), Inflammatory (C3), Lymphocyte Depleted (C4), Immunologically Quiet (C5) and TGF-β Dominant (C6).

### Mutational pre-processing

We used the R package dNdScv (version 0.0.1.0) from Martincorena et al to discover positively selected genes. We chose GRCh38 as our reference genome when using dNdScv. We ran dNdScv twice on our data: the first time to have the annotation from this package and the second time to obtain the selection results. Moreover, we performed an additional analysis during which we specified a list of driver genes (see driver_genes_im2017.txt in Data).

### Statistics and data visualisation

The ggplot2 (version 3.3.5) and ggpubr (version 0.4.0) packages were used for data visualisation. We divided patients into escape+ and escape-cohorts depending on whether they had a mutation in one genes from the immune list. Thus, our definition of immune evasion is from a genetic perspective. When reporting significantly selected genes, with the log of wmis being superior to 1, we decided to exclude olfactory receptors as they are spuriously mutated in cancer and unlikely to have a cancer driver role. For the confidence interval figures, we plotted the average global dNdS using mis_mle and we included the highest and lowest values of the cohort. Regarding the correlation analysis, we coloured the positively selected genes depending on whether they belonged to escape+, escape-escape or both.

### Survival

Using the library RTCGA.clinical (version 20151101.24.0) and the package survminer (version 0.4.9), we obtained the patient vital status at different time points. We divided patients into escaped and non-escaped cohorts to compare the survival between these two groups. This analysis was carried pancancer, on the 31 cancer types we are interested in and on the immune categories from Thorsson et al [17].

### Chi-square plots

To compare the distribution of escaped and non-escaped mutations and unique sites we used ggstatsplot::ggbarstats (version 0.9.1).

### Lolliplots

We used cBioPortal to create lolliplots showing the different locations and types of mutations between escaped and non-escaped cohorts pancancer. We focused on the driver genes of our pancancer analysis.

### Mutational signatures

To find the most dominant signatures, we used the deconstructSigs (version 1.8.0) R package. This enables us to find mutational signature associated to each cohort. We then looked at the most frequent signatures and reported the 11 dominant signatures which occurred most often. This methodology was applied to the pancancer cohort and various TCGA cancer types. We also plotted the proportion of mutations specific to the escape groups by running deconstructsigs on the escaped/nonescaped pancancer cohorts as a single sample.

## Notes

### Competing Interest Statement

The authors have declared no competing interest.

