## Supplementary figures and images for "Immune evasion impacts the selective landscape of driver genes during tumorigenesis"

### SF1

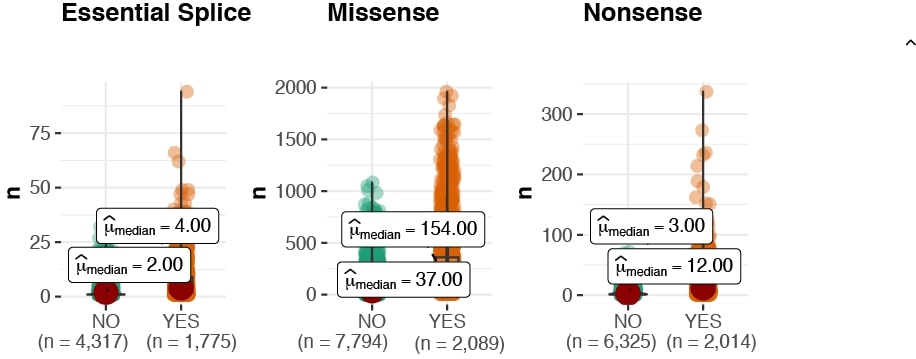

### SF2

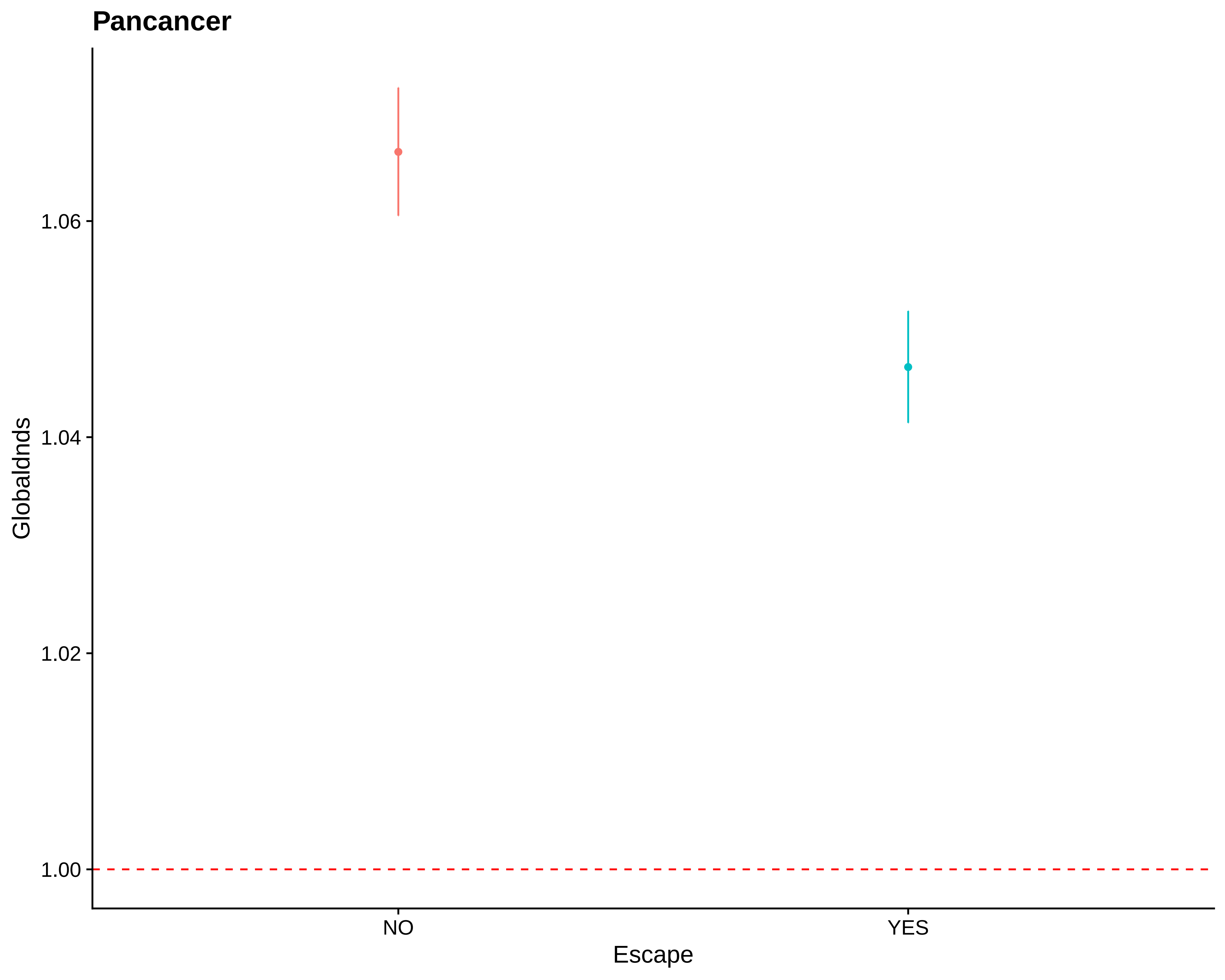

### SF3

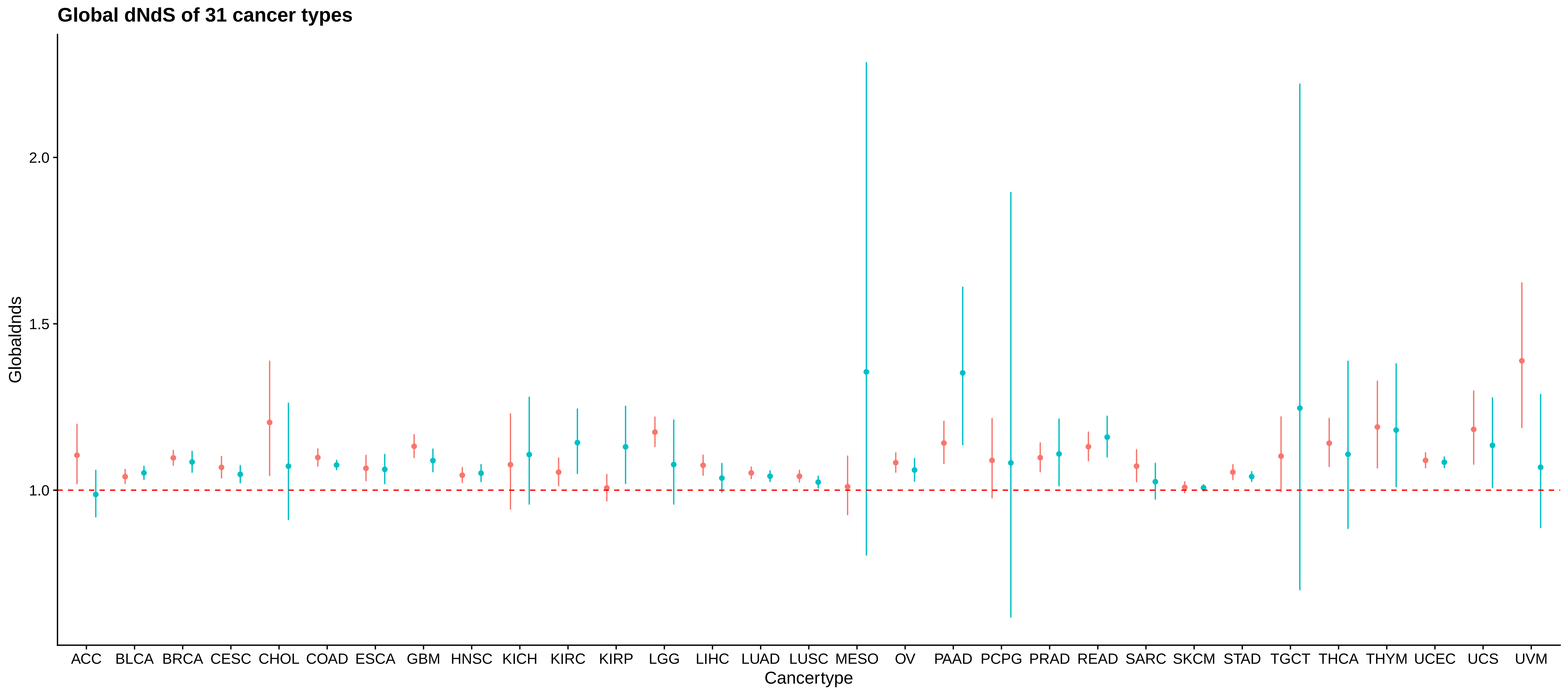

### SF4

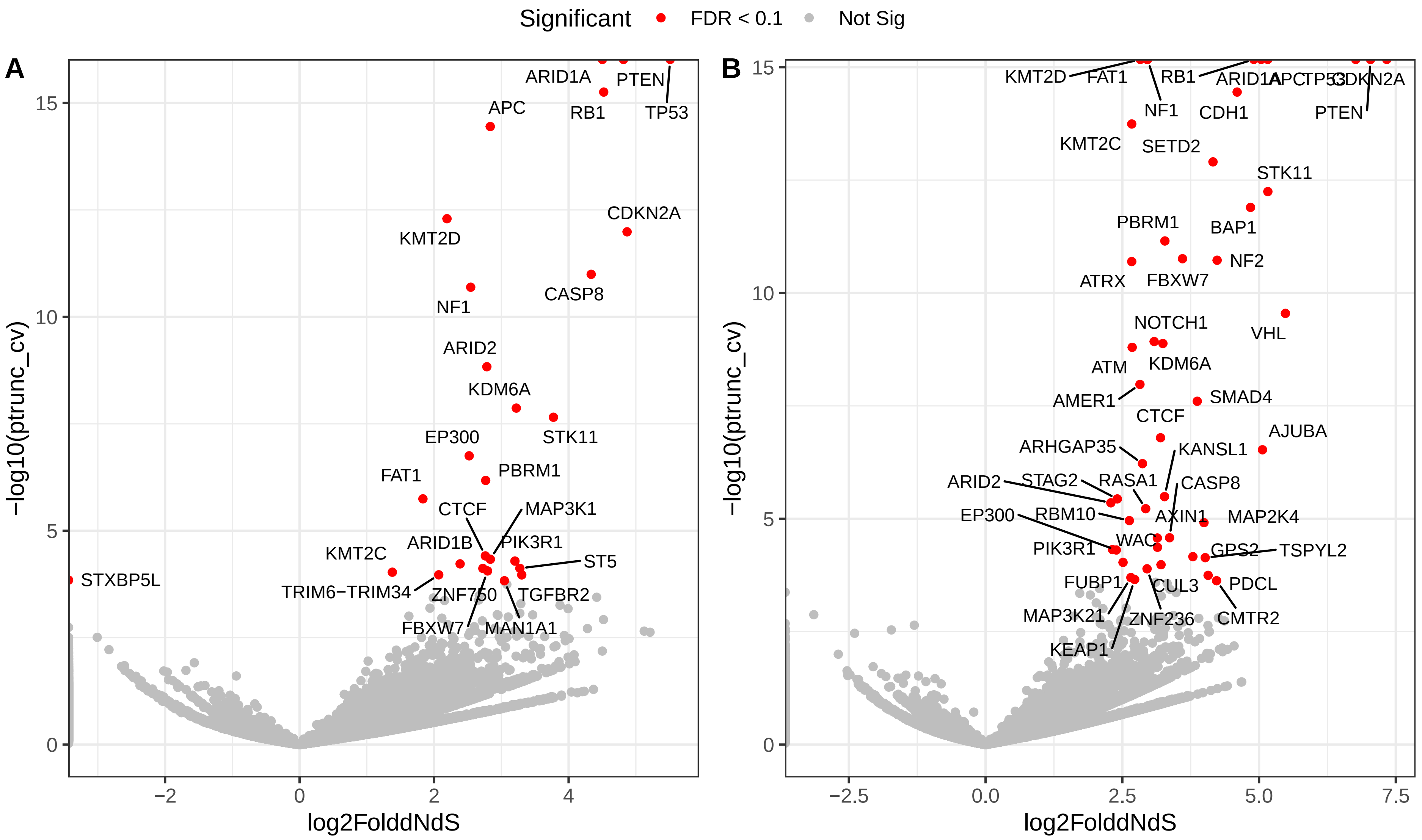

### SF5

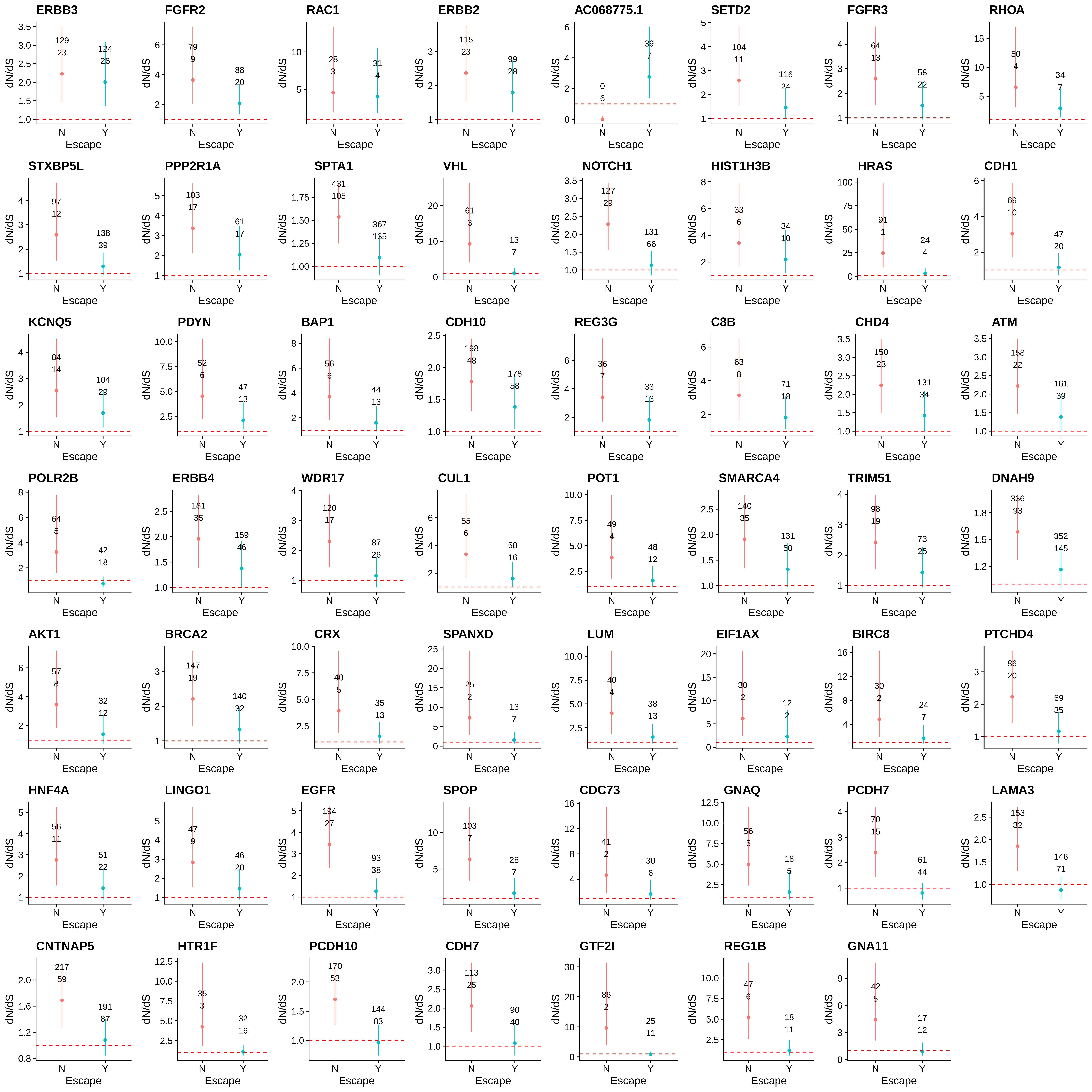

### SF6

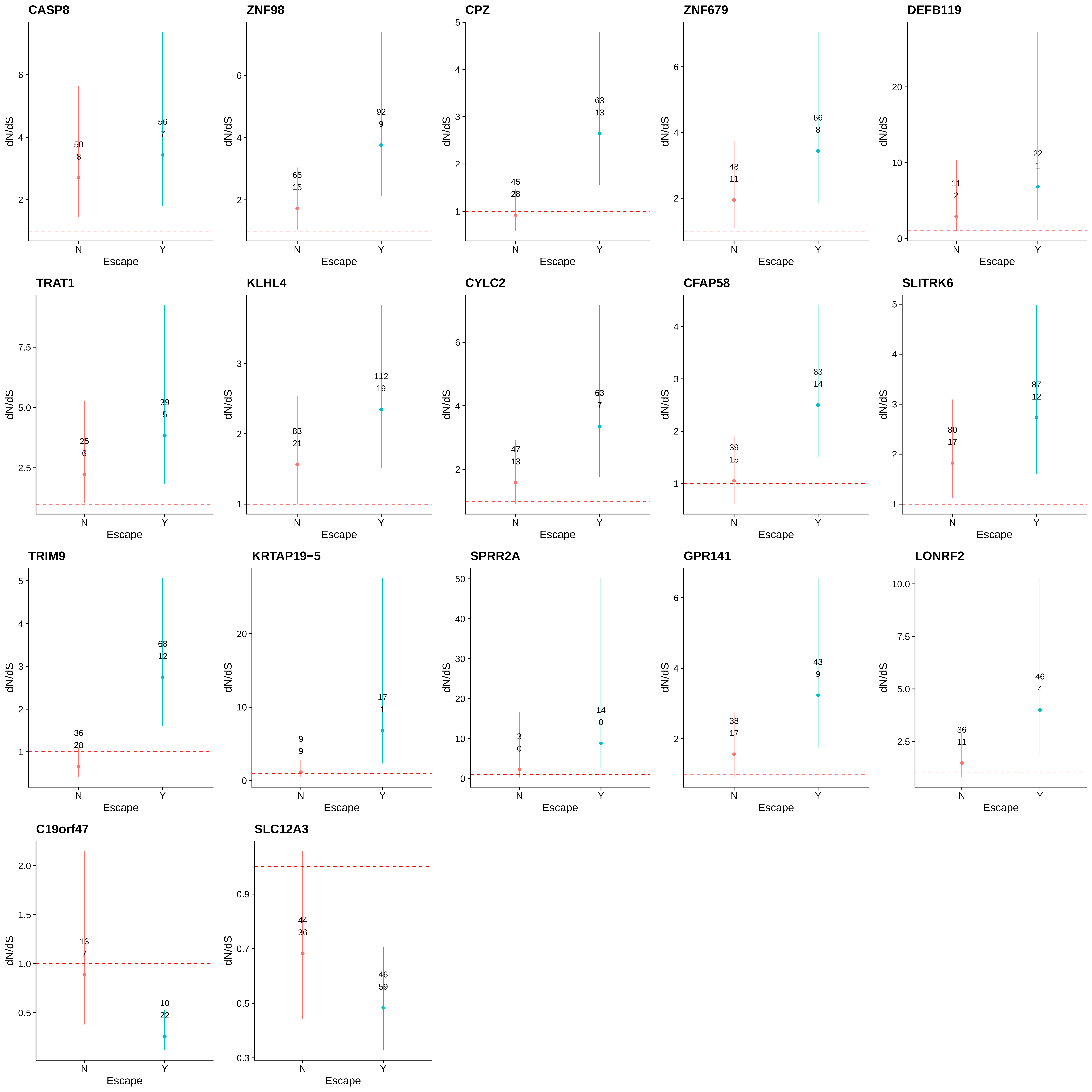

### SF7

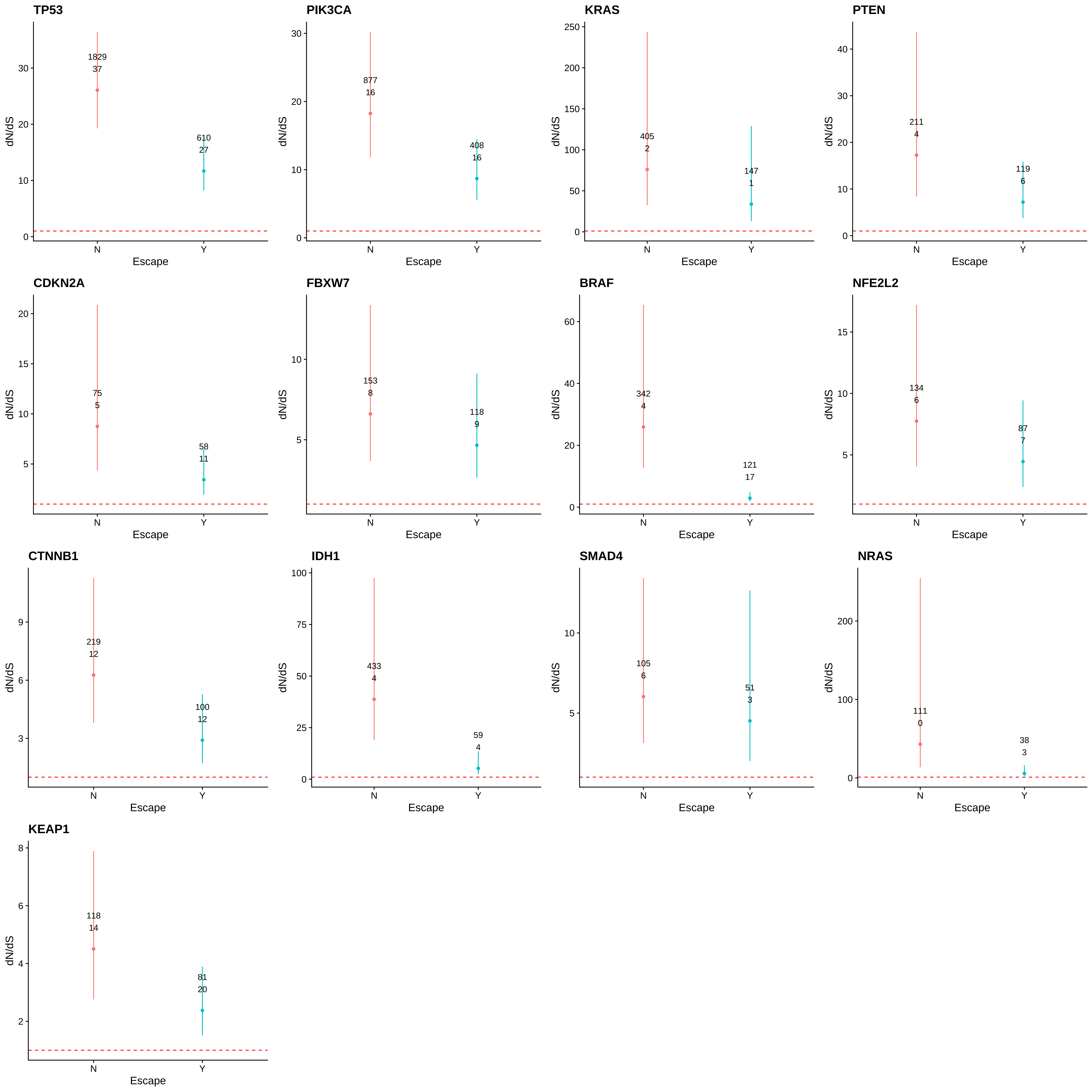
